# The influence of sensory noise, confidence judgments, and accuracy on pupil responses and associated behavioural adjustments in young and older people

**DOI:** 10.1101/2024.05.01.589279

**Authors:** Sara F. Oliveira, Catarina F. Gomes, Maria J. Ribeiro

## Abstract

We investigated the association between pupil responses, decision confidence and decision accuracy, their relationship with behavioural adjustments important for optimal task performance, and if these were altered in older adults.

We tested young and older adults, including men and women, with a dot motion direction discrimination task, while measuring task-related pupil responses. Feedback was provided after each trial. Participants were instructed to report perceived motion direction and simultaneously their decision confidence.

Older adults were overconfident in the presence of high sensory noise and in incorrect trials. Young people’s pupil responses reflected performance lapses. This effect was reduced in older people that showed blunted pupil responses to lapses and to negative feedback. Low confidence, errors, and larger pre-feedback pupil responses were associated with slower reaction time on the subsequent trial. The effects of errors and pupil were stronger in the older group.

In conclusion, older people’s blunted pupil responses following lapses suggest reduced awareness of their own abilities and internal brain state and might impair adjustment of behaviour for optimal performance.

## 1. **Introduction**

Metacognition is the ability to reflect on one’s own behaviour and can be assessed as the ability to identify erroneous responses without external feedback or the ability to assign confidence to onés own decisions in a way that predicts the correctness of the response (Fleming, 2024). Metacognitive awareness plays a role in perceptual learning (Guggenmos et al., 2016) and trial-by-trial behavioural adjustments that optimize performance (Desender et al., 2019).

Aging is associated with deficits in metacognition. Error awareness and metacognitive accuracy, the association between decision confidence and response accuracy, decrease with ageing (Harty et al., 2017; Overhoff et al., 2021; Palmer et al., 2014; Sim et al., 2020; Wessel et al., 2018). The ability to detect errors correlates with task performance suggesting that impaired metacognitive awareness affects older people’s behavioural output in decision-making tasks (Wessel et al., 2018). Interestingly, although deficits in metacognitive accuracy have been found in both memory (Dodson and Krueger, 2006; Martschuk et al., 2019; Souchay et al., 2007) and perceptual decision-making tasks (Klever et al., 2022; Palmer et al., 2014), the impairments appear more pronounced in perceptual tasks (Palmer et al., 2014). However, not all studies find an age-related deficit in metacognition. Specific task characteristics, differences in perceived task difficulty and delivery method might affect the findings (McWilliams et al., 2023).

The mechanism through which metacognitive awareness leads to behavioural adjustments, task learning and performance improvements might involve the engagement of pupil-linked arousal systems. Pupil dilation responses under isoluminant conditions follow the activation of neuromodulatory systems of the brainstem including noradrenergic and cholinergic systems (Joshi et al., 2016; Murphy et al., 2014; Reimer et al., 2016). These neuromodulators are important for learning and might guide behavioural adaptations associated with metacognitive awareness as well as explicit feedback processing (Crouse et al., 2020; de Leo et al., 2022). Low decision confidence is associated with higher engagement of pupil-linked arousal systems than high decision confidence (Allen et al., 2016; Colizoli et al., 2022; Urai et al., 2017). Pupil responses are also larger after errors than correct responses and larger after detected than undetected errors further supporting a link between confidence and the amplitude of the post-choice pupil dilation response (Critchley et al., 2005; Wessel et al., 2011). Pupil responses to feedback are associated with learning rate in reinforcement learning tasks (Van Slooten et al., 2018) and are modulated by confidence judgments (de Gee et al., 2021). Moreover, pupil responses following erroneous responses predict post-error slowing (Murphy et al., 2016) and older people show larger post-error slowing than young adults (Band and Kok, 2000; Ruitenberg et al., 2014; Wessel et al., 2018). Thus, age-related changes in the engagement of pupil-linked arousal systems might have an impact on behavioural adjustments important for optimal performance.

Pupil-linked arousal responses in perceptual decision-making tasks are mostly preserved in older people (Ribeiro and Castelo-Branco, 2019). In Ribeiro and Castelo-Branco (2019), the only difference across young and older adults in pupil responses during an auditory discrimination task was observed after the motor response where the pupil analyses suggested a less sustained response after the decision. This difference could reflect reduced activity in the arousal systems during the performance monitoring period. Accordingly, Wessel et al. (2018) showed that, in older people, pupil responses to errors are reduced and associated with reduced error awareness abilities (Wessel et al., 2018).

In the current study, we used a dot motion direction discrimination task to explore in further detail, in humans, how confidence, accuracy and sensory noise modulate post-decision pupil-linked arousal responses and trial-by-trial behavioural adjustments. To evaluate how these processes change with aging, we tested a group of young and a group of older adults. Reduced engagement of pupil-linked arousal in older people during the response monitoring interval might affect behavioural adaptation strategies important for task performance in perceptual decision-making tasks.

## 2. Materials and methods

### 2.1. Participants

Twenty-one healthy young adults and twenty-five healthy older adults were recruited for the study. All participants had normal or corrected-to-normal vision, and no current or previous history of vascular, neurological, or psychiatric disease. Exclusion criteria also included current use of psychoactive substances such as anxiolytics, antidepressants, antipsychotics, or beta-blockers, a history of chemotherapy, head trauma, and pregnancy. Participants were requested to abstain from alcohol consumption the night before. Nineteen older participants were evaluated with the “Montreal Cognitive Assessment – MoCA” (Nasreddine et al., 2005) and five with The Addenbrooke’s Cognitive Examination Revised (ACE-R) (Firmino et al., 2018; Mioshi et al., 2006) as screening measures to determine the global cognitive status. One older participant was not evaluated due to lack of availability. One older participant was excluded due to having a MoCA score more than 2SD below the mean expected for his age and education level, according to the normative data for the Portuguese population (Freitas et al., 2011). All other participants showed normal cognitive status. Three other older participants were excluded from analyses due to exhibiting very low task accuracies (around 50 %, chance level) even for the highest motion coherence stimulus. Therefore, the final sample for analyses included 21 young participants and 21 older participants. Table 1 shows the participants’ characteristics.

**Table 1.** Participants’ characteristics.

|  | N | Age (years) |  |  | Sex |  | Education (years) |  |  | Handedness |  |
| --- | --- | --- | --- | --- | --- | --- | --- | --- | --- | --- | --- |
|  |  | Mean | Min | Max | F | M | Mean | Min | Max | Left | Right |
| Young | 21 | 24 | 20 | 28 | 11 | 10 | 16 | 14 | 21 | 2 | 19 |
| Older | 21 | 60 | 50 | 72 | 11 | 10 | 16 | 10 | 25 | 1 | 20 |

The study was approved by the Ethics Committee of the Faculty of Medicine of the University of Coimbra (CE-162/2022). After explaining verbally and in writing the purpose and possible consequences of the experimental design, all participants signed the written informed consent.

### 2.2. Perceptual Decision-Making Task

We adapted a motion discrimination two-alternative choice task originally developed by Colizoli et al. (2018) (Colizoli et al., 2018). The task run in MATLAB (The MathWorks Company Ltd), with the Psychophysics Toolbox, version 3 (Brainard, 1997), and was presented on a computer monitor (19-inch Dell monitor) with a spatial resolution of 1440 x 1080 pixels and a refresh rate of 100 Hz. The monitor dimensions spanned 52.5 cm in width and 39.5 cm in height.

Dot motion stimuli were displayed within a central annulus with an outside diameter measuring 16.8° and an inner diameter of 2.4° with borders that were not visible to the participants. The fixation target was a white combination of bull’s eye and crosshair shape located in the centre of the annulus, as recommended by Thaler et al. (2013) for experiments that require stable fixation (Thaler et al., 2013). The dot stimuli were randomly distributed within the annulus and comprised two categories: signal dots, dots that moved coherently in either left or right directions at a speed of 7.5°/s, and noise dots, dots that moved at random (0 % coherence). Each frame contained a total of 524 white dots, each measuring 0.15° in diameter. The proportion of dots moving coherently vs. randomly defined the motion coherence, which modulates task difficulty. In the current work, motion coherence varied in four different levels (3, 6, 12, or 24 %).

Each trial was divided into five phases (Fig. 1): the baseline period (0.5–4.5 s); the stimulus interval consisting of random and coherent motion (0.75 s); the delay period preceding feedback (2.25 s); the feedback and intertrial interval (1.5–2.5 s). Except for the stimulus interval, random motion (0% coherence) was presented in all phases. Stimulus onset was indicated by an 880 Hz pure tone auditory cue with a duration of 250 ms. Feedback was also presented by an auditory signal consisting of the Portuguese words “Certo”, “Errado”, and “Não Respondeu”, respectively for correct, wrong, or no response. All auditory sounds were broadcast via a hi-fi speaker system at a volume clearly detectable by all participants. Both the central fixation target and the set of moving dots had a luminance of 58 cd/m² and were presented on a grey background, with a luminance of 43 cd/m². Participants were instructed to continuously maintain fixation on the central region and to indicate the perceived direction of coherent motion (left or right) and the confidence in their response by pressing a key on the computer keyboard. Participants should use the middle or index fingers of the left hand to indicate perception of leftwards motion and the middle or index fingers of the right hand to indicate perception of rightward motion, with the index fingers linked to low confidence responses and the middle fingers linked to high confidence responses, as represented in Figure 1.

**Figure 1.**
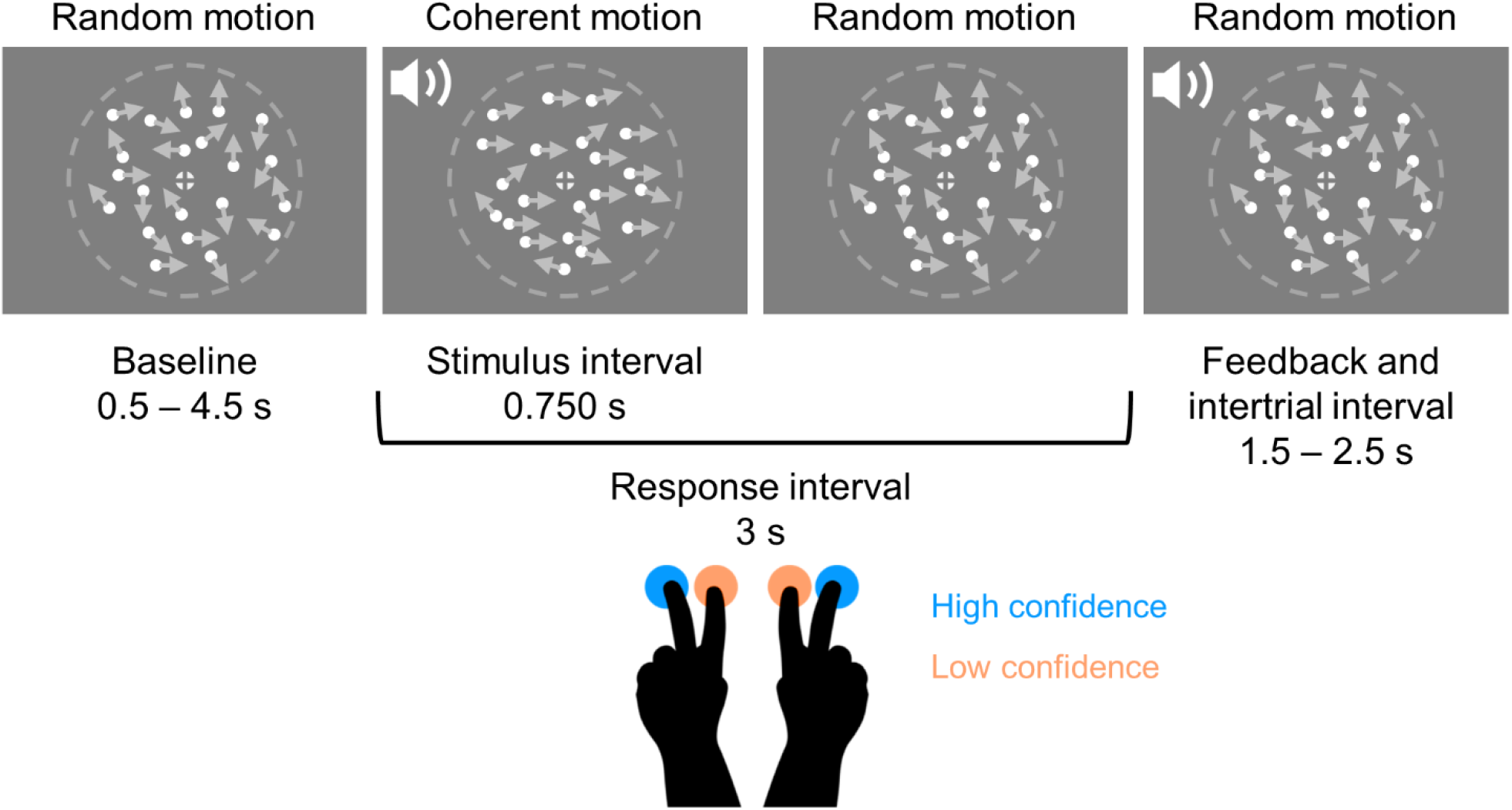
Schematic representation of the two alternative dot motion direction discrimination task. Participants were instructed to fixate the centre of the screen throughout the task. Trials started with a baseline period where randomly moving dots were presented. The start of the coherent motion stimulus was signalled by an auditory cue. Coherent motion was presented for 750 ms. Participants were instructed to respond by simultaneously indicating the perceived motion direction (left or right motion direction) and their confidence in the response (high confidence judgments were signalled by responding with left or right middle fingers and low confidence with left or right index fingers) within a 3 s window containing the motion stimulus. Auditory feedback was provided in every trial. Note that stimulus sizes in the panels are not to scale.

Before the experimental session, each participant completed a training session. The training consisted of the presentation of a trial with high coherence motion for each direction, followed by a sequence of three phases with eight high coherence motion trials, eight low coherence motion trials, and ten mixed high and low coherence trials, in which the stimulus could assume any of the coherence levels. During the training sessions, participants were instructed to indicate the perceived direction of coherent motion and the associated response confidence. The participants responses were monitored. If any difficulties were identified, the sequence of trials was repeated until the participants performed the task correctly.

Participants completed four runs of the task, lasting approximately eight minutes each, with self-paced breaks in between runs. The task consisted of 192 trials, with 48 trials per coherence level.

### 2.3. Behavioural Analyses

Reaction times were measured from stimulus onset to button press. For all analyses, we excluded the trials associated with impulsive responses where reaction time was lower than 100 ms.

### 2.4. Pupillometry Data Acquisition and Preprocessing

The EyeLink 1000 Plus desktop mount (SR Research, Ottawa, Ontario, Canada) was used to record horizontal and vertical gaze position and the pupilogram, monocularly at a sampling rate of 500 Hz. A trigger pulse was generated at stimulus onset, button press and feedback onset. Eye position calibration and validation was run before every run.

Pupil data was analysed with custom MATLAB scripts and the EEGLAB toolbox (Delorme and Makeig, 2004) in MATLAB R2022b. Missing data and blinks, as detected by the EyeLink software, were padded by 150 ms and linearly interpolated. Additional blinks were found using peak detection on the velocity of the pupil signal and linearly interpolated (Urai et al., 2017). Pupil data were normalized within each run to the percentage of the mean.

We analysed the pupil responses in the interval between the decision and the feedback in response-locked epochs with 200 ms baseline just before stimulus onset subtracted. For analyses of the pupil response after feedback, the epochs were locked with the onset of the feedback and the 200 ms baseline located just before feedback onset was subtracted. Epochs where more than half of the points in the interval from -1 s and 3 s from button press or feedback onset were interpolated were eliminated.

### 2.5. Statistical Analyses

Statistical analyses were conducted using IBM SPSS Statistics, version 27.0.1.0 software. Reaction time and pupil data were analysed by trial-wise linear mixed models (LMM). Generalized linear mixed models (GLMM) with logistic regression were used when the binary variables accuracy and confidence were the dependent variable. The degrees of freedom from these models represent the total number of trials minus the degrees taken up by the factors included in the model. Mixed models were chosen as these are robust to differences in the number of trials across conditions and missing data.

The factors of interest were considered fixed effects and the intersubject variability was considered the random effect. Only random intercepts were considered when building the models. The models were full factorial unless stated otherwise.

For the LMM analysis of pupil response and reaction time, the following factors of interest were included: group, motion coherence, confidence, and accuracy. As a control analysis, we also run the LMM of pupil response before feedback including as a covariate reaction time (main effect only) to control for the impact of reaction time on pupil size. For the GLMM analysis of accuracy, the following factors were included: group, motion coherence, and confidence. For the GLMM analysis of confidence, the following factors were included: group, coherence, and accuracy.

For the LMM analyses of the pupil responses, we included as dependent variable the mean amplitude within five 500 ms non-overlapping time windows from 500 ms up to 3000 ms after decision or feedback. So, for each analysis (pre and post feedback), we run five LMMs. To control for multiple comparisons, we considered significant only the effects with a *p* value less than or equal to 0.01 (Bonferroni correction).

To investigate the effect of trial confidence and accuracy on the reaction time of the subsequent trial, we used LMM analyses where we included as dependent variable the reaction time of the subsequent trial (trial n+1), and as factors motion coherence, confidence, and accuracy of trial n and group (main effects and interactions). We controlled for slow drifts in reaction time by including the main effect of the reaction time of the previous trial (trial n-1) (Murphy et al., 2016). For these analyses, we only included the trials where the trial n+1 and trial n-1 were correct trials.

To investigate the effect of the amplitude of the pupil responses on the reaction time of the subsequent trial, we used LMM analyses where we included as dependent variable the reaction time of the subsequent trial (trial n+1), and as factors pupil amplitude on trial n (one model for each of the five pupil 500 ms time windows) and group (main effects and interactions). We controlled for confidence and accuracy of trial n by including their main effects and for slow drifts in reaction time by including the main effect of the reaction time of the previous trial (trial n-1) (Murphy et al., 2016). For these analyses, we only included the trials where the trial n+1 and trial n-1 were correct trials.

Including motion coherence, confidence judgements and accuracy as predictors in the models posed the danger of multicollinearity as these variables were correlated. However, the correlation coefficients from Spearman correlation analyses were all lower than 0.310 suggesting that multicollinearity does not pose a threat for the interpretation of the findings.

## 3. Results

### 3.1. Older people respond to high sensory noise stimuli and in incorrect trials with higher confidence than young adults

Young and older adults were tested on a two alternative dot motion direction discrimination task and instructed to report motion direction and decision confidence simultaneously through button press (Fig. 1). Confidence judgments of both age groups exhibited the key signatures of decision confidence (Fleming, 2024; Sanders et al., 2016). Confidence increased with the strength of sensory evidence (motion coherence; Fig. 2A; Table 2), was higher for correct than incorrect responses (Fig. 2B) and showed an interaction between the strength of sensory evidence and accuracy, increasing with sensory evidence for correct responses but decreasing or increasing significantly less for incorrect responses (Fig. 2G), suggesting that both age groups were reporting their decision confidence as instructed.

**Figure 2.**
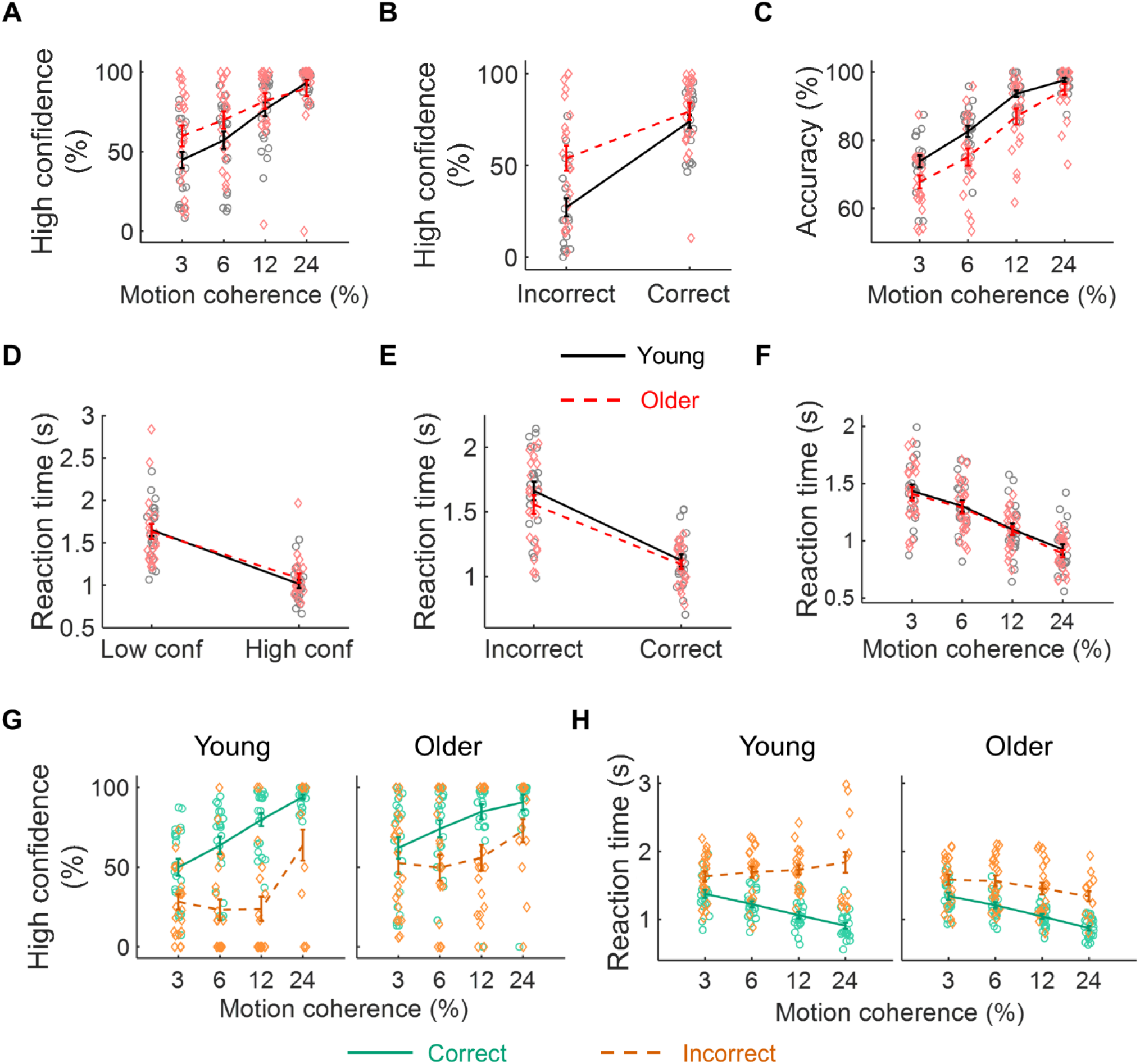
Behavioural performance of the older and young groups in the dot motion direction discrimination task. ***A***, Percentage of high confidence responses as a function of motion coherence. ***B***, Percentage of high confidence responses as a function of accuracy. ***C***, Accuracy as a function of motion coherence. ***D***, Reaction time as a function of response confidence. ***E***, Reaction time as a function of accuracy. ***F***, Reaction time as a function of motion coherence. ***A-F***, Black continuous line and grey circles represent data from the young group and the red dashed line and red diamonds represent data from the older group. Data are represented as mean ± standard error of the mean for each age group. ***G*** *and **H***, Percentage of high confidence responses (***G***) and reaction time (***H***) as a function of motion coherence for correct (green continuous lines) and incorrect (orange dashed lines) responses.

**Table 2.** Linear Mixed Models’ main effects and interactions with behavioural outputs, accuracy, confidence level or reaction time, as dependent variables.

|  | Accuracy |  | Confidence Level |  | Reaction Time |  |
| --- | --- | --- | --- | --- | --- | --- |
|  | *F* (*df*, *df*) | *p* | *F* (*df*, *df*) | *p* | *F* (*df*, *df*) | *p* |
| Group | **<u>14.3 (1, 7984)</u>** | **<u><.001</u>** | **<u>5.28 (1, 7984)</u>** | **<u>.022</u>** | 1.40 (1, 47) | .243 |
| Coherence | **<u>74.4 (3, 7984)</u>** | **<u><.001</u>** | **<u>54.5 (3, 7984)</u>** | **<u><.001</u>** | **<u>25.5 (3, 7960)</u>** | **<u><.001</u>** |
| Confidence | **<u>256 (1, 7984)</u>** | **<u><.001</u>** | --- | --- | **<u>343 (1, 7985)</u>** | **<u><.001</u>** |
| Accuracy | --- | --- | **<u>268 (1, 7984)</u>** | **<u><.001</u>** | **<u>348 (1, 7967)</u>** | **<u><.001</u>** |
| Group x Coherence | **<u>3.27 (3, 7984)</u>** | **<u>.020</u>** | **<u>3.02 (3, 7984)</u>** | **<u>.028</u>** | **<u>8.10 (3, 7960)</u>** | **<u><.001</u>** |
| Group x Confidence | **<u>22.7 (1, 7984)</u>** | **<u><.001</u>** | --- | --- | **<u>9.84 (1, 7985)</u>** | **<u>.002</u>** |
| Group x Accuracy | --- | --- | **<u>17.9 (1, 7984)</u>** | **<u><.001</u>** | 3.00 (1, 7967) | .083 |
| Coherence x Confidence | **<u>13.7 (3, 7984)</u>** | **<u><.001</u>** | --- | --- | 2.28 (3, 7960) | .077 |
| Coherence x Accuracy | --- | --- | **<u>16.7 (3, 7984)</u>** | **<u><.001</u>** | **<u>18.3 (3, 7961)</u>** | **<u><.001</u>** |
| Confidence x Accuracy | --- | --- | --- | --- | **<u>11.1 (1, 7966)</u>** | **<u><.001</u>** |
| Group x Coherence x Confidence | 1.36 (3, 7984) | .253 | --- | --- | .286 (3, 7960) | .836 |
| Group x Coherence x Accuracy | --- | --- | .724 (3, 7984) | .537 | 1.47 (3, 7961) | .222 |
| Group x Confidence x Accuracy | --- | --- | --- | --- | .203 (1, 7966) | .652 |
| Coherence x Confidence x Accuracy | --- | --- | --- | --- | .465 (3, 7960) | .707 |
| Group x Coherence x Confidence x Accuracy | --- | --- | --- | --- | .588 (3, 7960) | .623 |

Significant effects and interactions are underlined and highlighted in bold font.

The statistical results for the behavioural variables are reported in Table 2. Task accuracy was significantly reduced in the older group (accuracy mean ± standard deviation: young = 87 ± 5 %; older = 81 ± 8 %), yet older people responded more often with high confidence than young people (percentage of high confidence responses mean ± standard deviation: young = 68 ± 24 %; older = 74 ± 18 %). The effect of motion coherence on response confidence was weaker in older people that did not modulate their response confidence with the stimulus strength as much as young adults (significant group x coherence interaction; Fig. 2A). Older participants were more confident than young adults in incorrect responses but were equally confident in correct responses (significant group x accuracy interaction; Fig. 2B). For both groups of participants, response accuracy increased with increasing motion coherence (Fig. 2C). Finally, a significant group x coherence interaction reflected the fact that the effect of motion coherence on accuracy was stronger in the older group where accuracy was lower for the lower coherences but similar to the young group’s accuracy for the highest coherence.

Reaction time was associated with confidence judgments (Fig. 2D). In fact, the signatures of decision confidence were also evident in reaction time data, in both groups of participants. Reaction time was associated with accuracy (Fig. 2E) and motion coherence (Fig. 2F) and showed an interaction between sensory evidence and accuracy (Fig. 2H). The modulation of reaction time with confidence judgments was slightly weaker in the older group with a significant group x confidence interaction – the older group was slightly slower in the high confidence responses (Table 2). The effect of motion coherence on reaction time was slightly stronger in the older group that responded faster for the higher motion coherences (significant group x coherence interaction).

These results suggest that older participants were overconfident particularly when responding to stimuli with high sensory noise and in incorrect trials. In fact, seven older participants (33 %) responded with low confidence in less than 10% of the trials with one participant never responding with low confidence at all. All the young participants responded with low confidence in at least 10% of the trials.

### 3.2. Post-decision pupil response suggests blunted autonomic activation following performance lapses in the older group

The amplitude of the pupil-linked arousal responses between the decision and feedback (pre-feedback interval) were similar across age groups. However, significant interaction effects revealed specific pupil responses to lapses in young adults that were significantly reduced in older people.

Statistical results are described in Table 3. In the post-decision, pre-feedback interval, pupil dilation was higher for incorrect than correct responses. (Fig. 3A; Table 3). There was no significant effect of confidence judgements. However, we observed a significant confidence x accuracy interaction (Fig. 3B). This interaction was explored by running the LMM analyses separately for correct and incorrect trials (Supplemental Table 1). For correct responses, low decision confidence was associated with stronger pupil dilation than high confidence responses. In contrast, for incorrect responses, pupil responses to high confidence decisions (high confidence errors) were higher than pupil responses to low confidence errors. Thus, high confidence errors appear as an event where the autonomic system reacts as if the participant has very low confidence. A significant 3-way interaction group x confidence x accuracy suggests that this effect is blunted in the older group.

**Figure 3.**
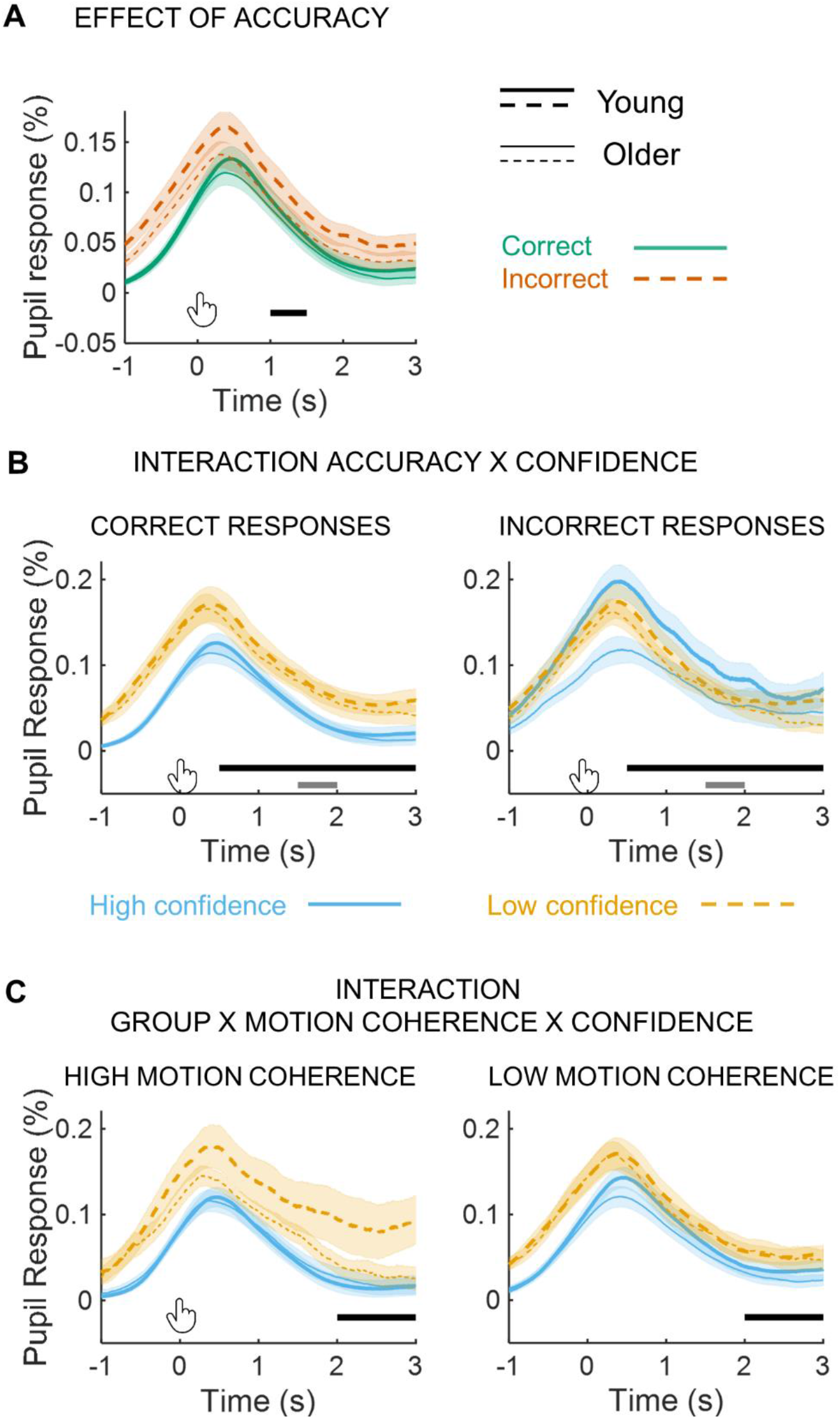
Task-related pupil responses locked with button press measured during the dot motion direction discrimination task. ***A***, Pupil responses as a function of response accuracy separately for the young and older groups. Orange dashed lines represent incorrect trials and green continuous lines represent correct trials. Horizontal black line indicates the time window where there is a significant effect of accuracy. ***B***, Pupil responses for correct (left) and incorrect (right) responses as a function of confidence judgments for both age groups. Horizontal black line indicates the time window where there is a significant confidence x accuracy interaction, while the horizontal grey lines indicate the time windows where there is a significant 3-way interaction group x confidence x accuracy. ***C***, Pupil responses for high motion coherence (left) and low motion coherence (right) as a function of response confidence for both age groups. Horizontal black line indicates time window where there is a significant 3-way interaction group x motion coherence x confidence. ***A****-**C***, Graphs represent mean ± standard error of the mean across participants. Thick lines represent the mean pupil responses of the young group and thin lines represent the mean pupil responses of the older group. ***B*** and ***C***, Yellow dashed lines represent pupil responses in low confidence trials and blue continuous lines represent pupil responses in high confidence trials.

**Table 3.** Linear mixed models’ main effects and interactions with the amplitude of the post-decision pupil response locked with button press averaged within each time window as dependent variable.

| Time window (s) | 0.5 - 1 |  | 1 - 1.5 |  | 1.5 - 2 |  | 2 - 2.5 |  | 2.5 - 3 |  |
| --- | --- | --- | --- | --- | --- | --- | --- | --- | --- | --- |
|  | *F* (df,df) | *p* | *F* (df,df) | *p* | *F* (df,df) | *p* | *F* (df,df) | *p* | *F* (df,df) | *p* |
| Group | 3.17 (1,48.8) | .081 | 2.8 (1,51.6) | .099 | 1.9 (1,55.9) | .176 | 2.1 (1,60.0) | .157 | 2.29 (1,64.80) | .135 |
| Coherence | 1.56 (3,7313) | .198 | 1.8 (3,7314) | .151 | 1.2 (3,7315) | .316 | .64 (3,7316) | .587 | .925 (3,7318) | .428 |
| Confidence | 2.82 (1,7339) | .093 | .71 (1,7346) | .400 | .03 (1,7352) | .872 | .03 (1,7351) | .865 | .109 (1,7346) | .741 |
| Accuracy | 5.39 (1,7321) | .020 | **<u>7.2 (1,7325)</u>** | **<u>.007</u>** | 6.2 (1,7331) | .012 | 4.3 (1,7335) | .038 | 2.51 (1,7339) | .113 |
| Group x Coherence | .618 (3,7313) | .603 | 9.4 (3,7314) | .419 | .63 (3,7315) | .595 | 1.1 (3,7316) | .358 | 1.99 (3,7318) | .113 |
| Group x Confidence | .034 (1,7339) | .854 | .39 (1,7346) | .535 | .93 (1,7352) | .335 | .59 (1,7351) | .442 | .022 (1,7346) | .882 |
| Group x Accuracy | 4.29 (1,7321) | .038 | 4.8 (1,7325) | .029 | 1.9 (1,7331) | .164 | 1.5 (1,7335) | .221 | .715 (1,7339) | .398 |
| Coherence x Confidence | 2.20 (3,7312) | .086 | 1.7 (3,7313) | .174 | 1.8 (3,7314) | .147 | 3.0 (3,7316) | .028 | 2.48 (3,7317) | .059 |
| Coherence x Accuracy | .809 (3,7313) | .489 | 1.2 (3,7314) | .321 | .87 (3,7316) | .455 | .43 (3,7317) | .733 | .394 (3,7319) | .757 |
| Confidence x Accuracy | **<u>11.6 (1,7319)</u>** | **<u><.001</u>** | **<u>9.2 (1,7322)</u>** | **<u>.002</u>** | **<u>12 (1,7327)</u>** | **<u>.001</u>** | **<u>13 (1,7330)</u>** | **<u><.001</u>** | **<u>9.01 (1,7334)</u>** | **<u>.003</u>** |
| Group x Coherence x Confidence | 1.75 (3,7312) | .154 | 1.7 (3,7313) | .173 | 2.7 (3,7314) | .042 | **<u>5.0 (3,7316)</u>** | **<u>.002</u>** | **<u>4.19 (3,7317)</u>** | **<u>.006</u>** |
| Group x Coherence x Accuracy | .264 (3,7313) | .852 | .45 (3,7314) | .716 | .64 (3,7316) | .600 | 1.4 (3,7317) | .231 | 1.27 (3,7319) | .284 |
| Group x Confidence x Accuracy | 3.30 (1,7319) | .069 | 5.2 (1,7322) | .022 | **<u>6.7 (1,7327)</u>** | **<u>.010</u>** | 4.4 (1,7330) | .035 | .787 (1,7334) | .375 |
| Coherence x Confidence x Accuracy | 1.50 (3,7312) | .212 | 1.7 (3,7313) | .168 | 3.2 (3,7314) | .021 | **<u>4.2 (3,7315)</u>** | **<u>.005</u>** | 3.15 (3,7317) | .024 |
| Group x Coherence x Confidence x Accuracy | 2.18 (3,7312) | .088 | 2.1 (3,7313) | .093 | 2.5 (3,7314) | .054 | **<u>3.9 (3,7315)</u>** | **<u>.009</u>** | 3.42 (3,7317) | .017 |

Significant effects and interactions after correction for multiple comparisons are underlined and highlighted in bold font.

Pupil response amplitude showed also a significant 3-way group x coherence x confidence interaction (Fig. 3C). For high motion coherence stimuli (low sensory noise), low confidence responses were associated with higher pupil responses than high confidence responses, and this effect is reduced for the low motion coherence stimuli (high sensory noise). In older people this effect was absent that is the effect of confidence on their pupil responses was not modulated by stimulus strength (Supplemental Table 2 shows the statistical results separately for the young and older groups). Therefore, in young but not in older adults, low confidence in response to stimuli that are easy to discriminate, possibly associated with periods of high internal noise, engage the autonomic system more than low confidence in response to stimuli that are difficult to discriminate.

Pupil responses are larger after longer reaction times and this effect might explain some of the results observed as low confidence and incorrect responses are associated with slower reactions. To investigate if the group differences observed in the amplitude of the pupil responses were associated with reaction time differences, we run the LMM analyses including reaction time as a covariate (Supplemental Table 3). All the effects were still significant, except the effect of accuracy.

### 3.3. Older people showed reduced effect of feedback valence on feedback evoked pupil responses

Feedback evoked a pupil response characterized by an initial pupil dilation followed by pupil constriction, as observed in previous studies (Chen et al., 2023; Van Slooten et al., 2018). The initial pupil dilation was modulated by feedback valence with negative feedback evoking higher pupil dilation responses than positive feedback (Fig. 4A; Table 4), while the later pupil constriction response was modulated by response confidence, with low confidence responses associated with a stronger pupil constriction (Fig. 4B; Table 4). Older people’s pupil responses showed a reduced effect of feedback valence with negative feedback evoking larger more sustained responses in the young group (significant group x accuracy interaction; Fig. 4A; Supplemental Table 4 shows statistical results separately for correct and incorrect trials). A significant confidence x accuracy interaction revealed that during the initial pupil dilation period, response confidence modulated the pupil response differently in correct versus incorrect trials (Fig. 4C; Supplemental Table 4). In the responses to positive feedback, pupil dilation was slightly higher for low confidence responses. In contrast, for negative feedback, the pupil responses were higher for high confidence responses (high confidence errors). The responses to high confidence errors appear reduced in the older group however the triple interaction group x confidence x accuracy was not significant after controlling for multiple comparisons (Table 4).

**Figure 4.**
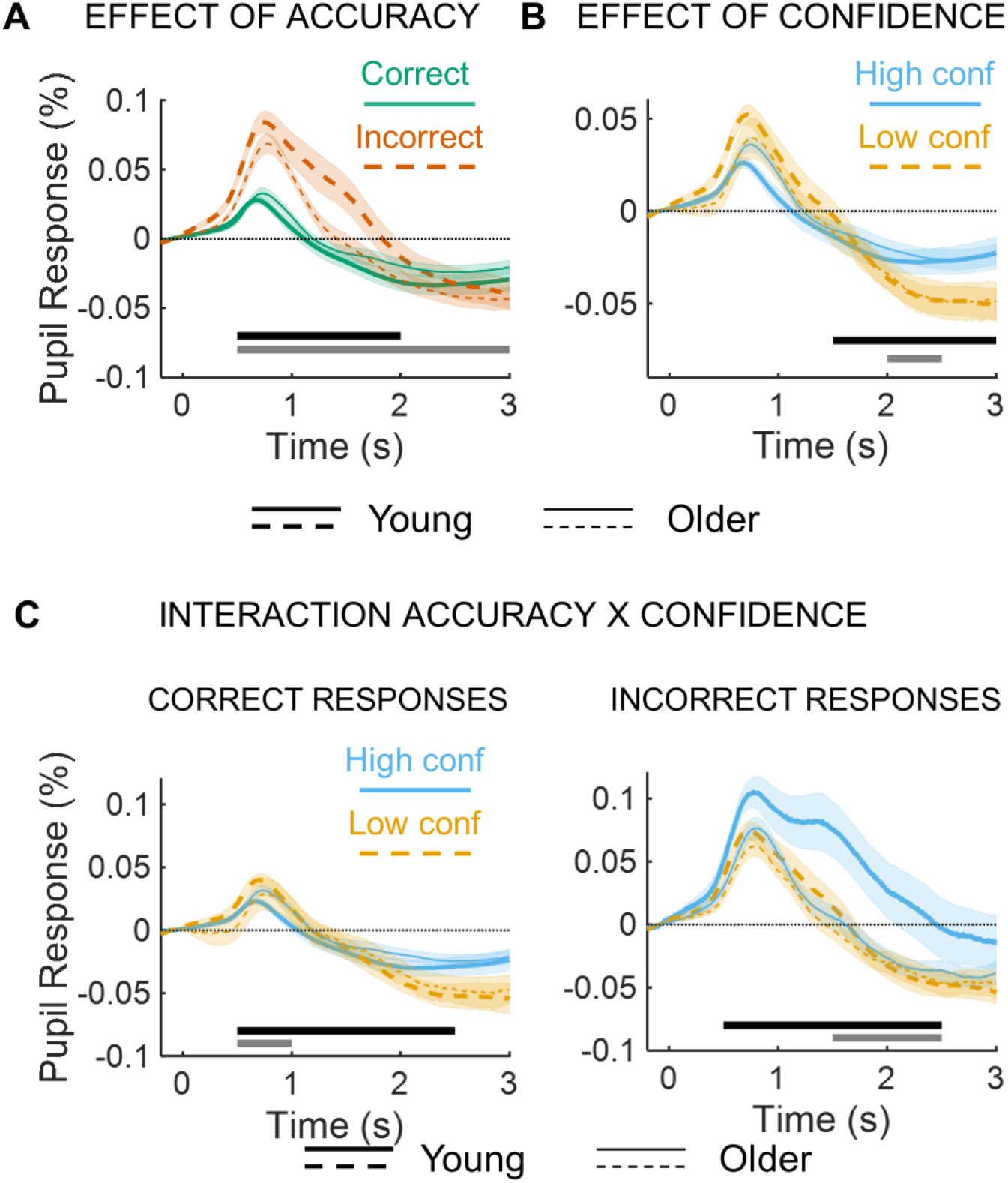
Task-related pupil responses locked with the auditory feedback. ***A***, Pupil responses as a function of response accuracy. Horizontal black line indicates the time window where there is a significant effect of accuracy, while the horizontal grey line indicates the time window where there is a significant group x accuracy interaction. ***B***, Pupil responses as a function of response confidence. Horizontal black line indicates the time window where there is a significant effect of confidence, while the horizontal grey line indicates the time window where there is a significant group x confidence interaction. ***C***, Pupil responses for correct (left) and incorrect (right) responses as a function of response confidence for both age groups. Horizontal black lines indicate the time window where there is a significant confidence x accuracy interaction, while the horizontal grey lines indicate the time windows where there is a significant effect of confidence when separating only correct or only incorrect trials. ***A***-***C***, Graphs represent mean ± standard error of the mean across participants. Thick lines represent the mean pupil responses of the young group and thin lines represent the mean pupil responses of the older group.

**Table 4.** Linear mixed models’ main effects and interactions with the amplitude of the pupil response locked with the auditory feedback averaged within each time window as dependent variable.

| Time window (s) | 0.5 - 1 |  | 1 – 1.5 |  | 1.5 – 2 |  | 2 – 2.5 |  | 2.5 - 3 |  |
| --- | --- | --- | --- | --- | --- | --- | --- | --- | --- | --- |
|  | *F* (df,df) | *p* | *F* (df,df) | *p* | *F* (df,df) | *p* | *F* (df,df) | *p* | *F* (df,df) | *p* |
| Group | 2.40 (1,84.8) | .125 | 4.4 (1,70.43) | .039 | 2.2 (1,72.50) | .140 | .40 (1,76.16) | .527 | .01 (1,69.73) | .913 |
| Coherence | 2.41 (3,7253) | .065 | 1.16 (3,7249) | .322 | .708 (3,7249) | .547 | .53 (3,7251) | .664 | .413 (3,7249) | .744 |
| Confidence | .010 (1,7173) | .921 | 4.09 (1,7249) | .043 | **<u>13.4 (1,7239)</u>** | **<u><.001</u>** | **<u>15.0 (1,7223)</u>** | **<u><.001</u>** | **<u>11.7 (1,7252)</u>** | **<u><.001</u>** |
| Accuracy | **<u>114 (1,7277)</u>** | **<u><.001</u>** | **<u>73.5 (1,7270)</u>** | **<u><.001</u>** | **<u>21.3 (1,7272)</u>** | **<u><.001</u>** | 1.37 (1,7273) | .243 | .652 (1,7270) | .419 |
| Group x Coherence | 1.66 (3,7253) | .173 | 1.99 (3,7249) | .113 | .929 (3,7249) | .426 | .348 (3,7250) | .790 | .175 (3,7259) | .914 |
| Group x Confidence | .623 (1,7173) | .430 | .889 (1,7249) | .346 | 3.57 (1,7239) | .059 | **<u>6.92 (1,7223)</u>** | **<u>.009</u>** | 6.03 (1,7252) | .014 |
| Group x Accuracy | **<u>15.7 (1,7277)</u>** | **<u><.001</u>** | **<u>34.6 (1,7270)</u>** | **<u><.001</u>** | **<u>30.3 (1,7272)</u>** | **<u><.001</u>** | **<u>13.8 (1,7273)</u>** | **<u><.001</u>** | **<u>7.13 (1,7270)</u>** | **<u>.008</u>** |
| Coherence x Confidence | 1.14 (3,7251) | .331 | 1.03 (3,7247) | .379 | .987 (3,7248) | .398 | 1.54 (3,7249) | .202 | 1.27 (3,7247) | .282 |
| Coherence x Accuracy | .824 (3,7254) | .480 | 1.09 (3,7249) | .353 | 1.07 (3,7249) | .359 | 1.23 (3,7251) | .296 | 2.20 (3,7249) | .086 |
| Confidence x Accuracy | **<u>14.2 (1,7272)</u>** | **<u><.001</u>** | **<u>13.0 (1,7264)</u>** | **<u><.001</u>** | **<u>15.6 (1,7266)</u>** | **<u><.001</u>** | **<u>6.94 (1,7168)</u>** | **<u>.008</u>** | 2.59 (1,7264) | .108 |
| Group x Coherence x Confidence | 1.07 (3,7251) | .362 | .449 (3,7247) | .718 | .893 (3,7248) | .444 | 1.95 (3,7149) | .119 | 2.21 (3,7247) | .084 |
| Group x Coherence x Accuracy | .809 (3,7254) | .489 | 1.66 (3,7249) | .174 | 1.20 (3,7250) | .308 | .968 (3,7251) | .407 | 1.76 (3,7249) | .152 |
| Group x Confidence x Accuracy | 2.22 (1,7272) | .136 | .856 (1,7264) | .355 | 2.48 (1,7266) | .116 | 4.31 (1,7168) | .038 | 2.02 (1,7264) | .155 |
| Coherence x Confidence x Accuracy | .710 (3,7251) | .546 | 1.34 (3,7247) | .260 | 1.31 (3,7248) | .269 | 1.74 (3,7148) | .156 | 1.17 (3,7247) | .321 |
| Group x Coherence x Confidence x Accuracy | 1.98 (3,7251) | .114 | 3.02 (3,7247) | .028 | 1.87 (3,7248) | .132 | 1.72 (3,7148) | .160 | 1.30 (3,7247) | .274 |

Significant effects and interactions after correction for multiple comparisons are underlined and highlighted in bold font.

### 3.4. Confidence, accuracy, and the amplitude of pre-feedback pupil response predict the reaction time of the subsequent trial

Next, we investigated the effect of accuracy and confidence on the reaction time of the subsequent trial. We found a significant effect of accuracy [reaction time was slower after errors than after correct trials; *F*_(1, 5572)_ = 15.9, *p* < .001; Fig. 5A], a significant effect of confidence [reaction time was slower after low confidence than after high confidence trials; *F*_(1, 5588)_ = 6.39, *p* = .011]. The effect of confidence did not depend on accuracy [no accuracy x confidence interaction; *F*_(1, 5566)_ = .139, *p* = .710] or on age group [no group x confidence interaction; *F*_(1, 5588)_ = .147, *p* = .701]. The effect of accuracy was larger in the older group than in the young group [group x accuracy interaction; *F*_(1, 5572)_ = 16.6, *p* < .001]. In fact, the effect of accuracy was only significant in the older group in linear mixed models with each group separate [young group *F*_(1, 2950)_ = .004, *p* = .950; older group *F*_(1, 2625)_ = 38.8, *p* < .001]. The effect of accuracy was confirmed with similar results using an approach adopted from the post-error slowing literature (Dutilh et al., 2012; see Supplementary Material). Interestingly, a group x coherence x confidence 3-way interaction was also observed [*F*_(1, 5557)_ = 3.03, *p* = .028], similar to the observed interaction in the post-decision pre-feedback pupil response. The effect of confidence depended on the stimulus motion coherence differently in young and older adults (Fig. 5B).

**Figure 5.**
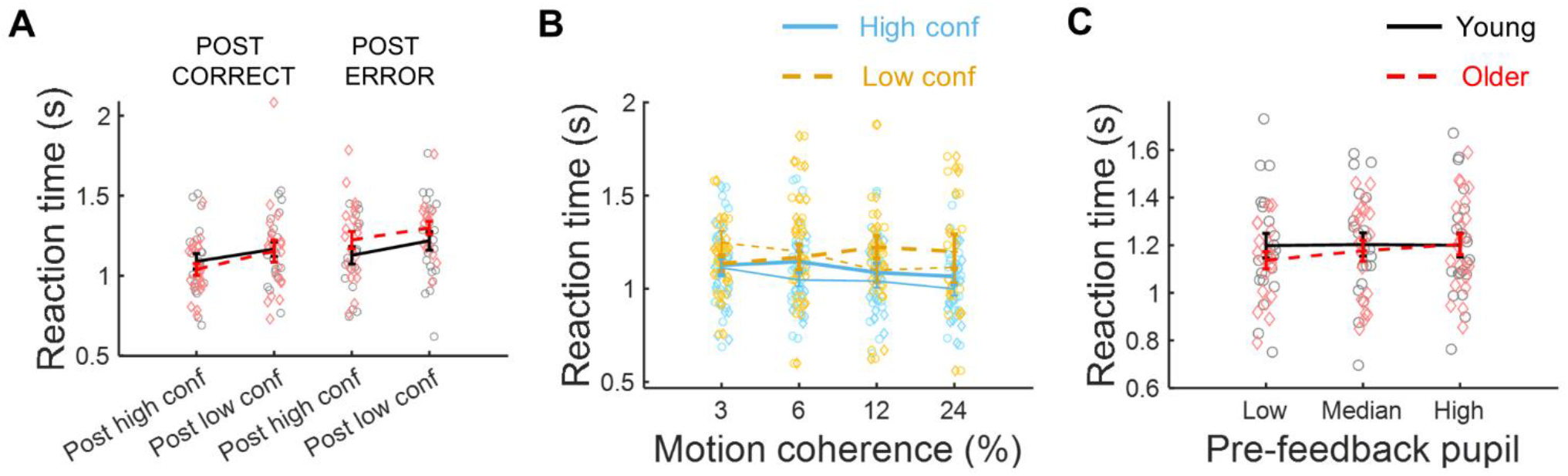
Effect of accuracy, confidence, motion coherence, and the amplitude of pre-feedback pupil responses on subsequent reaction time. ***A***, Reaction time sorted by confidence judgments and accuracy of the previous trial. ***B***, Reaction time sorted by confidence judgments and motion coherence of the previous trial. Thick lines and circle markers represent data from the young group and thin lines and diamond markers represent data from the older group. Continuous blue lines represent reaction time after high confidence decisions and orange dashed lines represent reaction time after low confidence decisions. ***C***, Reaction time sorted by the previous trial amplitude of the pupil response locked with button press within the time window between 2.5 and 3 s after button press. ***A*** and ***C,*** Black continuous line and grey circles represent data from the young group and the red dashed line and red diamonds represent data from the older group. **A-C**, Data are represented as mean ± standard error of the mean for each age group.

Finally, we investigated if the amplitude of the pupil responses before or after feedback predicted reaction time on the subsequent trial. The amplitude of the pupil responses before feedback in the time window between 2.5 and 3 s after button press was associated with reaction time on the subsequent trial [Fig. 5C; *F*_(1, 5229)_ = 7.38, *p* = .007]. This effect was larger in the older group [significant 2-way interaction group x pupil *F*_(1, 5119)_ = 6.82, *p* = .009 – young group only effect of pupil *F*_(1, 2708)_ = .008, *p* = .927; older group only *F*_(1, 2404)_ = 8.69, *p* = .003]. No other effect of pupil amplitude was observed (full statistical reporting in Supplemental Table 5).

## 4. Discussion

The amplitude of post-decision pupil-linked arousal responses before and after feedback was associated with decision confidence. These effects were observed in young and older adults. However, in comparison with young adults, older people showed reduced pupil dilation response after low confidence judgments to strong sensory evidence and after high confidence judgments in incorrect trials during the response monitoring period post-decision but before feedback, suggesting reduced autonomic modulation following performance lapses. Older people also showed reduced pupil responses to negative feedback. Trial-by-trial reaction time adjustments were modulated by decision confidence and accuracy. The effect of confidence was similar in young and older people, yet the effect of accuracy was larger in older people suggesting higher interference of errors on subsequent sensorimotor processing. Pre-feedback pupil response amplitude was associated with subsequent reaction time in the older group only, where larger pupil responses predicted subsequent slower responses.

### 4.1. Interactions between sensory noise, confidence judgments, and accuracy reveal reduced pupil responses to lapses in older people

As previously observed (Ribeiro and Castelo-Branco, 2019), the pupil responses from young and older adults evoked during perceptual decision-making were very similar with no significant group differences observed on the average responses. However, the effects of sensory noise, confidence judgments, and accuracy on the amplitude of the pupil responses by revealed interesting patterns that highlighted hidden cognitive processes that differ across age groups.

During the post-decision interval before feedback, low decision confidence in correct trials was associated with higher pupil dilation than high decision confidence. However, this pattern was inverted in incorrect trials where high decision confidence (high confidence errors) elicited a larger pupil response than low confidence errors. This suggests that high confidence errors are events of high cognitive conflict, maybe associated with lapses where the participants responded but immediately realized that the response was likely wrong. Interestingly, in semantic memory, high confidence errors are related to responses where the participants upon correction affirm that they knew the answer all along (Metcalfe and Finn, 2011). Activation of the autonomic system even before feedback might reflect this internal conflict. In contrast, older people do not show high pupil responses to high confidence errors as young people do suggesting lack of awareness of their own performance lapses, or different origin for high confident responses in error trials in this age group.

In line with the hypothesis that the pupil response to feedback reflects prediction errors (de Gee et al., 2021), we observed an interaction between confidence and accuracy after feedback. In the earlier phase of the pupil response to feedback, high confidence errors elicited the largest response while high confidence correct responses elicited the smallest response. Notably, this effect is already apparent before feedback suggesting a correlate of internally generated prediction errors. This effect both before and after feedback appears diminished in the older group where high confidence errors were not associated with larger pupil responses than low confidence errors. Studies on semantic memory suggest that high confidence errors are associated with a hypercorrection effect, where if participants are given the correct answers as feedback they are more likely to remember them on a retest than when they are given the answers after a low confidence error (Butterfield and Metcalfe, 2001) and this effect is impaired in older adults (Eich et al., 2013). The strong pupil-linked arousal response observed after feedback to a high confidence error might be important to facilitate memorization and explain why the effect is blunted in older people. In alternative, the large pupil responses observed in young people might reflect the internal conflict associated with the feeling of knowing the answer all along that might be reduced in older people.

Sensory noise also modulated the effect of confidence judgments on post-decision pupil responses. Low confidence responses to low sensory noise (high motion coherence) were associated with large pupil dilations during the post-decision interval before feedback. These responses were significantly diminished in older people. Low confidence responses to stimuli that should be easy to discriminate occur during periods where the percept has been corrupted by high internal noise, while low confidence in response to noisy stimuli can occur even during periods of low internal noise (Sanders et al., 2016). The fact that the pupil responses differ in these two conditions suggests that the arousal system is sensitive to the internal state of the perceptual system and distinguishes between internally generated noise and external noise. Periods of high internal noise might be associated with attention lapses for which humans can be immediately aware of (Arazi et al., 2019). The fact that older people’s pupils did not distinguish between conditions with high external or high internal noise suggests that ageing affects how the internal state modulates autonomic responses and might be linked to deficits in performance awareness. This is consistent with previous findings showing that older adults have impaired ability to detect their own performance errors (Harty et al., 2017) and show higher response confidence in unreported errors than young adults (Wessel et al., 2018).

### 4.2. Confidence and accuracy predict subsequent sensorimotor efficiency

Post-error slowing is thought to be a maladaptive response where error awareness interferes with the sensory processing of the subsequent stimulus leading to a compensatory slowdown of responses (Purcell and Kiani, 2016; Ullsperger and Danielmeier, 2016). Accordingly, post-error slowing is larger in children than adolescents or young adults (Smulders et al., 2016), larger in older people than young adults (Band and Kok, 2000; Ruitenberg et al., 2014; Wessel et al., 2018), and larger in people with lower cognitive abilities (Varriale et al., 2021).

We found that not only accuracy but also decision confidence influenced subsequent reaction time in an additive manner, in line with previous studies (Desender et al., 2019). Even after feedback, correct low decision confidence was associated with slower responses on the subsequent trial than correct high confidence responses, suggesting that this effect is related with the initial decision confidence and not objective trial accuracy. Interestingly, the effect of confidence was not modulated by accuracy suggesting that this effect is not associated with a prediction error signal where confidence in error trials should have the opposite effect of confidence in correct trials as observed in the pupil responses. That also suggests that pupil-linked arousal responses to feedback do not underlie post-error slowing as confirmed by our analyses where we did not observe a significant effect of the amplitude of the pupil response after feedback on subsequent reaction time. In fact, post-error slowing is observed regardless of feedback being given or not (e.g. Murphy et al., 2016; Purcell and Kiani, 2016; Wessel et al., 2018).

While the effect of confidence was observed in both age groups, the effect of accuracy was larger in the older group and not significant in the young group, suggesting higher error interference in the older group, as observed in previous studies (Band and Kok, 2000; Ruitenberg et al., 2014; Wessel et al., 2018).

Reaction time adjustments showed an interaction between group, motion coherence and confidence reflecting the fact that in the young group the effect of confidence was more pronounced in trials with high motion coherence, that is, low confidence in response to an easy to discriminate stimulus affected more the subsequent performance than low confidence in response to a difficult to discriminate stimulus. This observation parallels the observed effect on the post-decision pupil responses and suggests a behavioural consequence to the lack of arousal modulation in the older group in response to lapses caused by periods of high internal noise.

### 4.3. The impact of pupil-linked arousal on trial-by-trial behavioural adjustments

Brainstem ascending neuromodulatory systems associated with pupillary responses play a role in setting the brain state for optimal task performance modulating attentional focus, sensory processing, and task learning (Aston-Jones and Cohen, 2005; Murphy et al., 2021; Van Slooten et al., 2018). Post-error slowing has been shown to correlate with the amplitude of the pupil response post-error in a task where no feedback was presented (Murphy et al., 2016) and with the skin conductance response to errors (Hajcak et al., 2003) confirming an association with the sympathetic division of the autonomic nervous system. In our analyses, we observed an effect of pupil size in the pre-feedback period predicting subsequent reaction time only in the older group. It is important to note that in our task, we did not observe significant post-error slowing in the young group and this might have affected the ability to detect a link with the pupil responses. Thus, our findings confirm that in the group where post-error slowing was evident higher pupil dilation after the decision was associated with slower reaction times on the subsequent trial. However, the fact that the older group showed more pronounced post-error slowing and not larger pupil responses than the young group suggests the involvement of other mechanisms.

## 5. Conclusions

Decision confidence interacted with accuracy to predict pupil-linked arousal responses in a motion direction discrimination task during the post-decision pre-feedback period and after feedback in line with the idea that pupil responses are correlates of prediction error signals. Decision confidence and accuracy also modulated trial-by-trial reaction time adjustments albeit in an additive manner distinct from the prediction error processes suggesting that trial-by-trial behavioural adjustments reflect internal decision confidence independent of objective accuracy. Pupil responses suggested blunted autonomic responses to lapses in older people and reduced awareness of fluctuations in internal brain state also reflected in age-related differences in reaction time adjustments. Reduced autonomic activation following lapses might affect the ability of older people to quickly react to periods of inattention resulting in prolonged attention lapses. Post-error slowing was enhanced in the older group suggesting stronger error interference on subsequent sensorimotor processing that was also associated with pre-feedback pupil response.

## Data and code availability

Dataset publicly available at Open Science Foundation - https://osf.io/sq3eh/

MATLAB code for analyses available at GitHub - https://github.com/CIBIT-UC/2024_Oliveira_Confidence_pupil_aging

