## Supplementary Material for "The influence of sensory noise, confidence judgments, and accuracy on pupil responses and associated behavioural adjustments in young and older people"

**Supplemental Table 1.** Linear mixed models' main effects and interactions with the amplitude of the post-decision pupil response locked with button press averaged within each time window as dependent variable, separately for correct and incorrect trials.

| Time window (s) | 0.5 - 1 | | 1 – 1.5 | | 1.5 – 2 | | 2 – 2.5 | | 2.5 - 3 | |
|---|---|---|---|---|---|---|---|---|---|---|
| | *F* (df,df) | *p* | *F* (df,df) | *p* | *F* (df,df) | *p* | *F* (df,df) | *p* | *F* (df,df) | *p* |
| **Correct trials only** | | | | | | | | | | |
| Group | 1.39 (1,43.98) | .245 | .815 (1,44.86) | .372 | .619 (1,46.0) | .436 | .799 (1,47.0) | .376 | 1.39 (1,48.3) | .245 |
| Coherence | **3.85 (3,6153)** | **.009** | 3.59 (3,6157) | .013 | 3.72 (3,6161) | .011 | 3.56 (3,6164) | .014 | **4.3 (3,6167)** | **.005** |
| Confidence | **42.9 (1,6180)** | **<.001** | **26.2 (1,6146)** | **<.001** | **23.9 (1,6062)** | **<.001** | **21.4 (1,5956)** | **<.001** | **18 (1,5817)** | **<.001** |
| Group x Coherence | .545 (3,6153) | .652 | 1.01 (3,6157) | .388 | 1.55 (3,6161) | .199 | 2.48 (3,6164) | .059 | 2.6 (3,6167) | .053 |
| Group x Confidence | 4.38 (1,6180) | .036 | 3.53 (1,6146) | .060 | 3.55 (1,6062) | .060 | 2.67 (1,5956) | .102 | 2.0 (1,5817) | .162 |
| Coherence x Confidence | **5.66 (3,6154)** | **<.001** | 2.58 (3,6157) | .052 | 2.50 (3,6162) | .057 | 2.98 (3,6165) | .030 | 3.6 (3,6168) | .014 |
| Group x Coherence x Confidence | .833 (3,6154) | .475 | .685 (3,6157) | .561 | 1.10 (3,6162) | .350 | 1.27 (3,6165) | .283 | 1.3 (3,6168) | .281 |
| **Incorrect trials only** | | | | | | | | | | |
| Group | 4.57 (1,56.66) | .037 | 4.40 (1,58.80) | .040 | 3.34 (1,67.7) | .072 | 3.34 (1,75.89) | .072 | 3.15 (1,80.9) | .080 |
| Coherence | .943 (3,1135) | .420 | 1.62 (3,1136) | .182 | 1.15 (3,1141) | .326 | .356 (3,1145) | .785 | .31 (3,1147) | .821 |
| Confidence | 1.16 (1,1164) | .283 | 1.79 (1,1166) | .181 | 4.63 (1,1164) | .032 | **6.86 (1,1150)** | **.009** | 4.5 (1,1137) | .033 |
| Group x Coherence | .612 (3,1135) | .607 | .960 (3,1136) | .411 | 1.01 (3,1141) | .388 | 1.68 (3,1145) | .170 | 2.0 (3,1147) | .116 |
| Group x Confidence | 5.13 (1,1164) | .024 | 5.77 (1,1166) | .016 | **7.02 (1,1164** | **.008** | 4.43 (1,1150) | .036 | .80 (1,1137) | .372 |
| Coherence x Confidence | 1.65 (3,1134) | .176 | 2.22 (3,1135) | .084 | 3.72 (3,1139) | .011 | **5.31 (3,1143)** | **.001** | **3.8 (3,1145)** | **.010** |
| Group x Coherence x Confidence | 2.16 (3,1134) | .091 | 2.51 (3,1135) | .057 | **3.81 (3,1139)** | **.010** | **6.22 (3,1143)** | **<.001** | **5.0 (3,1145)** | **.002** |

Significant effects and interactions after correction for multiple comparisons are underlined and highlighted in bold font.

**Supplemental Table 2.** Linear mixed models' main effects and interactions with the amplitude of the post-decision pupil response locked with button press averaged within each time window as dependent variable, separately for the young and older groups.

| Time window (s) | 0.5 - 1 | | 1 – 1.5 | | 1.5 – 2 | | 2 – 2.5 | | 2.5 - 3 | |
|---|---|---|---|---|---|---|---|---|---|---|
| | *F* (df,df) | *p* | *F* (df,df) | *p* | *F* (df,df) | *p* | *F* (df,df) | *p* | *F* (df,df) | *p* |
| **Young group only** | | | | | | | | | | |
| Coherence | .855 (3,3677) | .464 | 1.18 (3,3678) | .316 | .621 (3,3679) | .602 | .492 (3,3679) | .688 | .848 (3,3680) | .468 |
| Confidence | .622 (1,3688) | .430 | .011 (1,3691) | .917 | .207 (1,3696) | .649 | .280 (1,3697) | .596 | .058 (1,3697) | .809 |
| Accuracy | 5.45 (1,3681) | .020 | **6.74 (1,3682)** | **.009** | 4.33 (1,3686) | .037 | 3.15 (1,3688) | .076 | 1.70 (1,3689) | .192 |
| Coherence x Confidence | 2.02 (3,3677) | .109 | 1.61 (3,3678) | .185 | 2.22 (3,3678) | .084 | **3.99 (3,3679)** | **.008** | 3.38 (3,3679) | .017 |
| Coherence x Accuracy | .393 (3,3677) | .758 | .739 (3,3677) | .529 | .481 (3,3678) | .695 | .814 (3,3679) | .486 | .742 (3,3679) | .527 |
| Confidence x Accuracy | **7.69 (1,3680)** | **.006** | **8.00 (1,3681)** | **.005** | **10.5 (1,3683)** | **.001** | **9.59 (1,3685)** | **.002** | 4.32 (1,3686) | .038 |
| Coherence x Confidence x Accuracy | 1.53 (3,3677) | .204 | 1.83 (3,3677) | .139 | 2.85 (3,3678) | .036 | **3.96 (3,3679)** | **.008** | 3.05 (3,3679) | .028 |
| **Older group only** | | | | | | | | | | |
| Coherence | 2.15 (3,3635) | .091 | 2.14 (3,3636) | .093 | 2.22 (3,3636) | .084 | 2.35 (3,3637) | .071 | **3.67 (3,3638)** | **.012** |
| Confidence | 3.93 (1,3649) | .047 | 2.43 (1,3652) | .119 | 1.49 (1,3654) | .222 | .477 (1,3655) | .490 | .069 (1,3651) | .793 |
| Accuracy | .084 (1,3639) | .772 | .312 (1,3642) | .576 | 1.54 (1,3643) | .215 | .868 (1,3645) | .352 | .631 (1,3648) | .427 |
| Coherence x Confidence | .370 (3,3635) | .775 | .338 (3,3635) | .798 | .260 (3,3636) | .854 | .487 (3,3636) | .691 | .479 (3,3637) | .697 |
| Coherence x Accuracy | .868 (3,3636) | .457 | .743 (3,3636) | .527 | 1.30 (3,3637) | .272 | .823 (3,3637) | .481 | .807 (3,3639) | .490 |
| Confidence x Accuracy | 3.05 (1,3638) | .081 | .664 (1,3640) | .415 | .966 (1,3641) | .326 | 2.97 (1,3643) | .085 | 5.36 (1,3634) | .021 |
| Coherence x Confidence x Accuracy | 1.22 (3,3635) | .299 | .216 (3,3635) | .885 | .310 (3,3636) | .818 | .445 (3,3636) | .721 | 1.12 (3,3637) | .338 |

Significant effects and interactions after correction for multiple comparisons are underlined and highlighted in bold font.

**Supplemental Table 3.** Linear mixed models' main effects and interactions with the amplitude of the post-decision pupil response locked with button press averaged within each time window as dependent variable, including reaction time and a covariate.

| Time window (s) | 0.5 - 1 | | 1 – 1.5 | | 1.5 – 2 | | 2 – 2.5 | | 2.5 - 3 | |
|---|---|---|---|---|---|---|---|---|---|---|
| | *F* (df,df) | *p* | *F* (df,df) | *p* | *F* (df,df) | *p* | *F* (df,df) | *p* | *F* (df,df) | *p* |
| Group | 2.82 (1,48.8) | .100 | 2.6 (1,51.9) | .117 | .36 (1,56.8) | .207 | 1.8 (1,50.43) | .191 | 1.89 (1,55.01) | .174 |
| Coherence | 1.96 (3,7313) | .119 | 2.1 (3,7314) | .095 | 1.2 (3,7316) | .227 | .67 (3,6202) | .574 | .864 (3,6204) | .459 |
| Confidence | .095 (1,7341) | .758 | .11 (1,7349) | .745 | 1.1 (1,7352) | .317 | 2.1 (1,6216) | .143 | 1.60 (1,6190) | .206 |
| Accuracy | .932 (1,7322) | .334 | 2.3 (1,7328) | .130 | 1.4 (1,7334) | .183 | .61 (1,6224) | .435 | <.001 (1,6228) | .982 |
| Group x Coherence | .308 (3,7312) | .820 | .59 (3,7314) | .624 | .52 (3,7315) | .738 | .93 (3,6201) | .424 | 1.70 (3,6203) | .164 |
| Group x Confidence | .185 (1,7339) | .667 | .69 (1,7347) | .405 | 2.2 (1,7352) | .239 | 1.0 (1,6219) | .316 | .016 (1,6196) | .901 |
| Group x Accuracy | 3.51 (1,7320) | .061 | 4.1 (1,7325) | .044 | 1.3 (1,7331) | .223 | 1.0 (1,6220) | .302 | .369 (1,6224) | .543 |
| Coherence x Confidence | 2.56 (3,7312) | .053 | 1.9 (3,7313) | .127 | 1.4 (3,7314) | .119 | 3.3 (3,6201) | .019 | 2.88 (3,6202) | .034 |
| Coherence x Accuracy | .839 (3,7312) | .472 | .99 (3,7314) | .398 | .39 (3,7316) | .663 | .22 (3,6203) | .882 | .189 (3,6205) | .904 |
| Confidence x Accuracy | **9.89 (1,7318)** | **.002** | **7.9 (1,7322)** | **.005** | **8.2 (1,7327)** | **.001** | **11 (1,6218)** | **.001** | **7.33 (1,6222)** | **.007** |
| Group x Coherence x Confidence | 1.65 (3,7312) | .175 | 1.6 (3,7313) | .184 | 2.3 (3,7315) | .044 | **4.9 (3,6201)** | **.002** | **4.07 (3,6203)** | **.007** |
| Group x Coherence x Accuracy | .191 (3,7312) | .902 | .37 (3,7314) | .777 | .62 (3,7316) | .605 | 1.5 (3,6202) | .215 | 1.29 (3,6204) | .275 |
| Group x Confidence x Accuracy | 3.36 (1,7318) | .067 | 5.3 (1,7322) | .022 | 5.5 (1,7327) | .009 | 4.5 (1,6218) | .033 | .834 (1,6222) | .361 |
| Coherence x Confidence x Accuracy | 1.54 (3,7312) | .201 | 1.8 (3,7313) | .154 | 2.9 (3,7314) | .019 | **4.4 (3,6201)** | **.004** | 3.27 (3,6202) | .020 |
| Group x Coherence x Confidence x Accuracy | 2.11 (3,7312) | .096 | 2.1 (3,7313) | .103 | 1.8 (3,7314) | .060 | **3.8 (3,6201)** | **.010** | 3.34 (3,6202) | .019 |
| Reaction time | **47.6 (1,7351)** | **<.001** | **35 (1,7339)** | **<.001** | **29 (1,7282)** | **<.001** | **44 (1,6074)** | **<.001** | **66.2 (1,5930)** | **<.001** |

Significant effects and interactions after correction for multiple comparisons are highlighted in bold font and underlined.

**Supplemental Table 4.** Linear mixed models' main effects and interactions with the amplitude of the pupil response locked with feedback averaged within each time window as dependent variable, separately for correct and incorrect trials.

| Time window (s) | 0.5 - 1 | | 1 – 1.5 | | 1.5 – 2 | | 2 – 2.5 | | 2.5 - 3 | |
|---|---|---|---|---|---|---|---|---|---|---|
| | *F* (df,df) | *p* | *F* (df,df) | *p* | *F* (df,df) | *p* | *F* (df,df) | *p* | *F* (df,df) | *p* |
| **Correct trials only** | | | | | | | | | | |
| Group | 1.41 (1,55.7) | .240 | 1.60 (1,52.5) | .211 | 2.96 (1,51.2) | .091 | 2.52 (1,52.05) | .118 | 1.87 (1,50.73) | .177 |
| Coherence | **4.55 (3,6141)** | **.003** | .688 (3,6141) | .559 | .714 (3,6141) | .543 | 2.31 (3,6141) | .074 | 2.14 (3,6140) | .092 |
| Confidence | **20.8 (1,4601)** | **<.001** | 2.16 (1,5227) | .141 | .042 (1,5346) | .838 | 4.40 (1,5208) | .036 | **2.46 (1,5453)** | **.006** |
| Group x Coherence | .286 (3,6141) | .836 | .748 (3,6141) | .523 | .159 (3,6141) | .924 | .231 (3,6141) | .875 | 1.09 (3,6140) | .352 |
| Group x Confidence | .206 (1,4501) | .650 | .026 (1,5227) | .871 | .323 (1,5346) | .570 | .468 (1,5208) | .494 | 1.45 (1,5453) | .228 |
| Coherence x Confidence | 1.09 (3,6141) | .352 | 1.07 (3,6142) | .359 | 1.52 (3,6141) | .206 | 1.53 (3,6142) | .205 | 1.19 (3,6141) | .312 |
| Group x Coherence x Confidence | .422 (3,6141) | .737 | 2.25 (3,6142) | .081 | 1.71 (3,6141) | .162 | .860 (3,6142) | .461 | 1.38 (3,6141) | .247 |
| **Incorrect trials only** | | | | | | | | | | |
| Group | 5.81 (1,63.5) | .019 | **7.48 (1,59.12)** | **.008** | 6.31 (1,67.3) | .014 | 1.75 (1,77.24) | .190 | .229 (1,74.21) | .634 |
| Coherence | .790 (3,1101) | .499 | .848 (3,1098) | .468 | .764 (3,1105) | .515 | .721 (3,1109) | .539 | 1.82 (3,1107) | .142 |
| Confidence | 4.74 (1,1123) | .030 | 5.23 (1,1126) | .022 | **9.56 (1,1103)** | **.002** | **7.47 (1,1087)** | **.006** | 3.91 (1,1107) | .048 |
| Group x Coherence | 1.70 (3,1101) | .167 | 1.77 (3,1098) | .150 | 1.21 (3,1105) | .303 | .779 (3,1109) | .506 | 1.24 (3,1107) | .295 |
| Group x Confidence | 1.07 (1,1123) | .300 | 1.33 (1,1126) | .248 | 3.75 (1,1103) | .053 | **6.78 (1,1087)** | **.009** | 4.85 (1,1107) | .028 |
| Coherence x Confidence | .917 (3,1098) | .432 | 1.36 (3,1096) | .254 | 1.59 (3,1101) | .191 | 2.19 (3,1105) | .088 | 1.70 (3,1104) | .165 |
| Group x Coherence x Confidence | 1.86 (3,1098) | .135 | 1.66 (3,1096) | .175 | 1.73 (3,1101) | .159 | 2.44 (3,1105) | .063 | 2.32 (3,1104) | .073 |

Significant effects and interactions after correction for multiple comparisons are underlined and highlighted in bold font.

**Supplemental Table 5.** Linear mixed models' main effects and interactions with reaction time on the following trial as dependent variable.

| Time window (s) | 0.5 - 1 | | 1 – 1.5 | | 1.5 – 2 | | 2 – 2.5 | | 2.5 - 3 | |
|---|---|---|---|---|---|---|---|---|---|---|
| | *F* (df,df) | *p* | *F* (df,df) | *p* | *F* (df,df) | *p* | *F* (df,df) | *p* | *F* (df,df) | *p* |
| **Pupil before feedback** | | | | | | | | | | |
| Group | .635 (1,48.1) | .429 | .770 (1,43.8) | .385 | .850 (1,42.1) | .362 | .811 (1,41.7) | .373 | .816 (1,41.7) | .372 |
| Pupil | .006 (1,5033) | .939 | .91 (1,5102) | .763 | 2.57 (1,5117) | .109 | 3.99 (1,5121) | .046 | <u>**7.38 (1,5119)**</u> | <u>**.007**</u> |
| Group x Pupil | .002 (1,5033) | .962 | .22 (1,5102) | .638 | 1.88 (1,5117) | .170 | 3.98 (1,5121) | .046 | <u>**6.82 (1,5118)**</u> | <u>**.009**</u> |
| **Pupil after feedback** | | | | | | | | | | |
| Group | 1.21 (1,42.3) | .278 | .843 (1,41.4) | .364 | .795 (1,41.6) | .378 | .810 (1,41.7) | .373 | .861 (1,41.7) | .359 |
| Pupil | .266 (1,5088) | .606 | .11 (1,5089) | .737 | .018 (1,5096) | .895 | .022 (1,5093) | .882 | .152 (1,5096) | .696 |
| Group x Pupil | 3.63 (1,5090) | .057 | 1.4 (1,5088) | .237 | .290 (1,5096) | .590 | .035 (1,5094) | .851 | .088 (1,5095) | .767 |

Significant effects and interactions after correction for multiple comparisons are highlighted in bold font and underlined.

**Robust post error slowing analysis**

*Methods*

Post error slowing was calculated by comparing reaction time on correct trials that followed error trials with reaction time on correct pre-error trials that followed a correct trial (Dutilh et al., 2012). This method ensures equal number of post error and post correct trials equally distributed throughout the task thus controlling for possible drifts in response accuracy with time-on-task. To account for interindividual differences in overall reaction time, post error slowing was quantified in percent change from post correct reaction time (Wessel et al., 2018).

Post error slowing was compared across groups using independent samples *t*-test and one-sample *t*-test analyses in MATLAB (The MathWorks Company Ltd).

*Results*

As expected, reaction time after errors was slower than reaction time after correct trials (reaction time mean ± SD: young post correct = 1.141 ± 0.225 s; young post error = 1.168 ± 0.241 s; older post correct = 1.096 ± 0.177 s; older post error = 1.201 ± 0.169 s). Post error slowing was stronger in older people, however, this difference did not reach statistical significance [post error slowing mean ± SD: young = 3.10 ± 15.5 %; older = 10.4 ± 11.0 %; $t_{(40)} = -1.76$, $p = .086$]. Nevertheless, when testing each group separately, post error slowing was only significantly different from zero in the older group [one-sample *t*-test: young $t_{(20)} = .918$, $p = .370$; older $t_{(20)} = 4.33$, $p < .001$].
